# Luminance and contrast of images in the THINGS database

**DOI:** 10.1101/2021.07.08.451706

**Authors:** William J Harrison

## Abstract

The THINGS database is a freely available stimulus set that has the potential to facilitate the generation of theory that bridges multiple areas within cognitive neuroscience. The database consists of 26,107 high quality digital photos that are sorted into 1,854 concepts. While a valuable resource, relatively few technical details relevant to the design of studies in cognitive neuroscience have been described. We present an analysis of two key low-level properties of THINGS images, luminance and luminance contrast. These image statistics are known to influence common physiological and neural correlates of perceptual and cognitive processes. In general, we found that the distributions of luminance and contrast are in close agreement with the statistics of natural images reported previously. However, we found that image concepts are separable in their luminance and contrast: we show that luminance and contrast alone are sufficient to classify images into their concepts with above chance accuracy. We describe how these factors may confound studies using the THINGS images, and suggest simple controls that can be implemented a priori or post-hoc. We discuss the importance of using such natural images as stimuli in psychological research.

## Introduction

In recent decades, advances in computational power and analytical methods have resulted in a growing emphasis on collecting, curating, and sharing large datasets in all areas of science. So called “big data” provides an opportunity to uncover relatively high-order structure through exploratory analyses, experimentation, or a combination of both (e.g. Hebart et al., 2020). The THINGS database is an image database of 26,107 high quality digital photos that have been sorted into 1,854 labelled “concepts” (Hebart et al., 2019). This database is a massive stimulus set that has the potential to advance understanding of perceptual and cognitive processes associated with more complex stimuli than typically used in lab-based settings (Carandini et al., 2005). The THINGS images are freely available (http://doi.org/10.17605/osf.io/jum2f), but relatively few technical details relevant to the design of studies in cognitive neuroscience have been described. The aim of the present report is thus to provide a summary of variation in luminance and luminance contrast within the THINGS database. We anticipate that these data will inform and constrain the design of studies in which luminance and contrast are expected to correlate with dependent variables of primary interest.

A great deal of visual neuroscientific theory has been developed from studies that vary the luminance and luminance contrast properties of stimuli. Indeed, there is a rich history of describing the responses of individual visual neurons as a function of contrast (e.g. Hubel & Wiesel, 1959). Much is also known about the ways in which luminance and contrast influence perception of abstract stimuli (e.g. Campbell & Robson, 1968) and natural images (e.g. Geisler, 2008). However, the purpose of this brief summary is not to provide an in-depth review of the role of basic image properties in perception and cognition – this information is available in most psychology textbooks as well as many review papers (Graham, 1989; Pelli & Bex, 2013). Instead, we emphasise that it is well understood that the most fundamental of visual processes are highly contingent on changes in luminance and contrast. In many experimental designs, therefore, it is important to control for these image attributes if they are not of interest to the researcher.

The THINGS database has already been leveraged to advance understanding of perception and cognition. By combining human judgements of similarity with computational modelling, Hebart et al (2020) quantified the psychological dimensions on which natural images vary. Their model describes 49 attributes that capture variations in conceptual and perceptual properties of the THINGS images. We recently used the THINGS images to investigate how a set of image statistics contribute to observers’ ability to detect targets in natural images (Rideaux et al., 2021). In both studies, variability in low-level image properties are directly or indirectly related to the researchers’ hypotheses. However, unquantified contrast differences across THINGS images may confound experimental manipulations in other cases. Grootswagers et al (2021) recently made publicly available the THINGS-EEG database, which includes the electroencephalography (EEG) responses of 50 participants to 22,248 THINGS images, with 12 images from each of all 1,854 concepts in the database. EEG responses are known to be influenced by image contrast (Campbell & Maffei, 1970), and so analyses of this dataset may require careful treatment of the metrics we report. A further aim of the present report, therefore, is to investigate whether luminance and contrast differences across THINGS images are sufficient for above-chance decoding of image concepts.

Differences in luminance and luminance contrast across ten randomly sampled THINGS images can be seen in Figure 1. Although the THINGS images are full colour (24-bit; top row), we quantify achromatic variations only, as shown by the greyscale images in the middle row. The bottom row shows the log amplitude of the Fourier transformed images. The differences in distributions of contrast energy are somewhat unsurprising, given that the amplitude simply re-expresses the image (albeit in a phase-invariance format). The analyses described below systematically quantify 1) general patterns of luminance and contrast differences across the images and concepts, and 2) the degree to which concepts can be predicted from item-level differences in the luminance and contrast of THINGS images.

**Figure 1.**
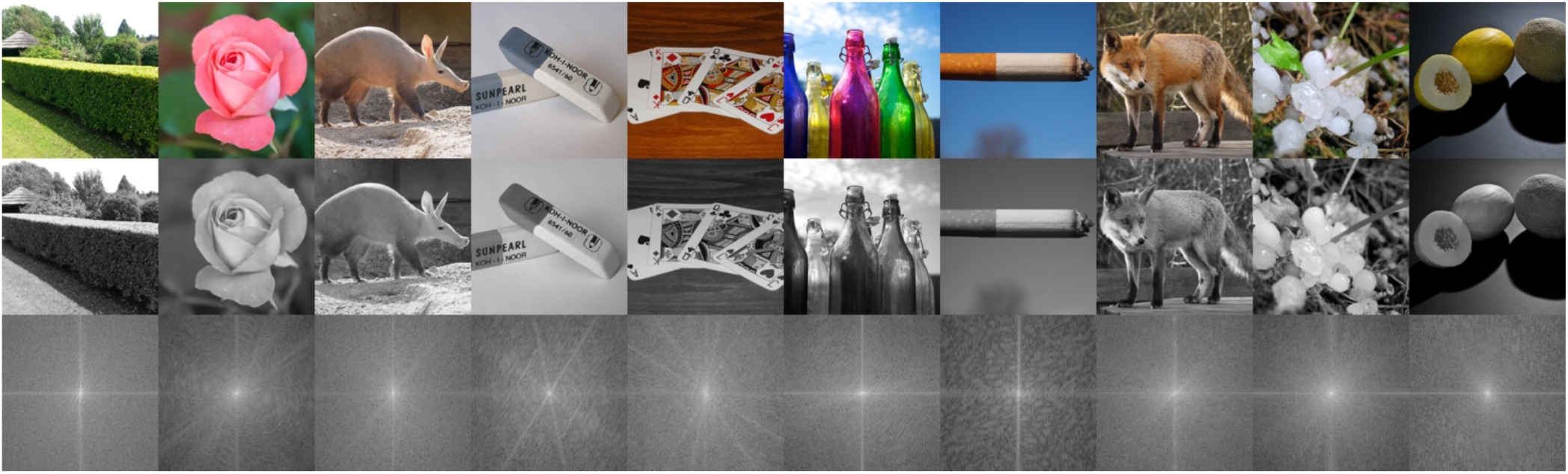
Example images from the THINGS database. Top row: images from 10 randomly chosen concepts. Middle row: greyscale versions of the example images. Bottom row: log amplitude of the Fourier-transformed images, rescaled between 0 (black) and 1 (white).

## Methods

All data and code are shared via the Open Science Framework: https://osf.io/v8a3q/. The THINGS database can be accessed via: http://doi.org/10.17605/osf.io/jum2f.

### Image pre-processing

We converted each image in the THINGS database to greyscale using MATLAB’s (MathWorks) rgb2gray() function, converted the image from a uint8 to double, and divided all values by 255 to initially set the range of the image to fall between 0 and 1. Images were rescaled to 512 × 512 pixels using Matlab’s imresize() function using bicubic interpolation. The minimum image size for inclusion in the original THINGS dataset was 480 × 480 pixels, but most images were 600 × 600 or larger. Therefore, resizing to 512 × 512 resulted in a size decrease for the vast majority of images. We selected this size because it is smaller than the majority of THINGS images while being an integer power of two, the combination of which make some of our image processing analyses more efficient in the frequency domain. We discuss this issue further in the Discussion. Due to pixel interpolation, some values of the resized images fell below or above 0 and 1, respectively. Such values were clipped.

Digital images are typically encoded with a compressive nonlinearity to over-represent relatively darker tones, and we assumed the same is true of the THINGS images (in a non-exhaustive search, we did not find any images that included the colour profile in the metadata). We therefore linearised the images prior to measuring their image spectra via gamma-correction:

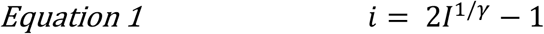

Where *I* is a source image in the range 0 – 1, *γ* is the assumed encoding gamma, and *i* is the linearised image. We set *γ* to 2 based on the findings of Bex et al (2005) who showed that this value provides comparable amplitude spectra as correctly calibrated images. The other terms in Equation 1 scale the image from -1 to 1, which is convenient for interpreting the following measures. Error in the estimate of *γ* should be inconsequential to the perceptual appearance of images (Bex et al., 2005), but will result in small errors in the luminance calculations performed here; we are not aware of a better alternative without ground truth knowledge of each image’s encoding function. Varying *γ* is unlikely to affect the general patterns of data or conclusions we report. Perhaps more important is that an experimenter selects a consistent method when measuring images and for the correct linearised display of images in the lab.

### Luminance

Luminance is a measure of the intensity of light from a given area of space after the light has passed through a model of the sensitivity of a standard eye. Although environmental luminance can be approximated from calibrated digital images using metadata regarding the camera’s encoding functions (e.g. Frazor & Geisler, 2006), we do not have access to such information for the THINGS database. When considering the use of these images for lab-based experiments, however, the critical luminance values are those of the digital image, and the luminance of pixels on a computer display is perfectly correlated with the pixel values. With the image in the range -1 to 1 (see *Image pre-processing*, above), we express relative luminance as varying linearly from the darkest black possible tone on a given monitor, to mid-grey, to the most luminous tone possible (i.e. values -1, 0, and 1, respectively). Mean luminance, therefore, is simply the mean of an image’s pixel values:

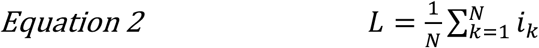

Where *k* is a pixel index, *N* is the total number of pixels, and *L* is the mean luminance.

### RMS contrast

Luminance contrast can be measured in different ways. Bex and Makous (2002) found that root-mean-square (RMS) contrast was the best predictor of observers’ sensitivity to natural images. RMS contrast (*C*_*rms*_) is simply the standard deviation of the luminance (i.e. pixel) values:

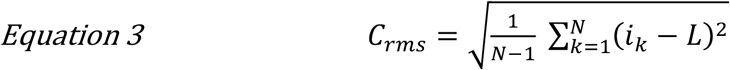

There are other methods with which to quantify contrast, such as finding the difference between the maximum and minimum luminance values, or finding the difference between the maximum absolute value and the average value. For simplicity, however, we quantify only RMS contrast across the full image. We also use other measures of contrast to quantify energy in separable spatial frequency and orientation bands, as we describe below.

### Correlation between luminance and RMS contrast

We quantified the relationship between luminance and RMS contrast using a generalised linear multilevel model (GLMM) to predict contrast from luminance. The model can be written as:

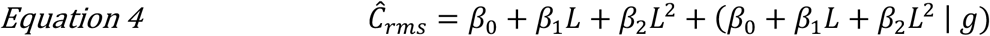

Where *β*_*n*_ are the beta weights, *g* is the grouping factor (image concept), and *Ĉ*_*rms*_ is the estimated RMS contrast. We used a multilevel model because it uses the grouping of images into concepts to improve the estimate of the relationship between luminance and contrast: it quantifies the relationship between luminance and contrast for each concept simultaneously, while also diminishing the influence of concepts that are relatively highly variable in these measures (Gelman & Hill, 2007). We fit the model using Matlab’s fitglme() function.

### Spatial frequency filtered contrast

An image’s contrast energy at any given spatial frequency (and/or orientation) can be quantified according to amplitude in the frequency domain (Figure 1, bottom row). To compute contrast energy as a function of spatial frequency, therefore, we simply grouped the Fourier-transformed amplitude values according to their distances from the origin and found the mean of each group. Fourier amplitude is calculated as the absolute of the Fourier-transformed complex values. Prior to analysis of the Fourier amplitude, images were windowed inside a circular mask (radius = 256 pixels) with a cosine edge, smoothly transitioning the edges of the image to mid-grey over 12 pixels. The Fourier amplitude was windowed in the same aperture to remove off-cardinal information that had a greater frequency than the maximum frequency of cardinal orientations (i.e. maximum frequency = 256 cycles/image).

Our 512 × 512 images cover 8 octaves (1 – 256 cycles/image). To calculate contrast energy in evenly spaced spatial frequency bands, we first computed the distance of each point in the amplitude spectrum from the origin based on the x/y distances using Pythagoras’s theorem. We then log_2_-scaled these values and applied the floor function to round-down each log-scaled distance to an integer. The resulting values express the distance of each point from the origin according to 1-octave wide spatial frequency bands. We then averaged the amplitude values within each band.

### Orientation filtered contrast

To quantify variations in contrast energy according to the orientation of the contrast, we summed the Fourier amplitude within each of 16 oriented bands that ranged the full circle (0 ± 90°). Oriented filters were raised cosine filters that ramped from 0 to 1 (or 1 to 0) over 11.25°. The filters therefore had a bandwidth of 22.5° at the base, and 11.25° at half-height, and extended across all spatial frequencies. Filters were evenly spaced in 11.25° steps, such that summing across all filters resulted in a value of 1 at all locations (i.e. the filters were a “complete” basis set).

### Modelling spatial frequency and orientation contrast energy

We fit models that summarise the distributions of contrast energy across spatial frequencies and orientations. Contrast energy across spatial frequencies was estimated with the function:

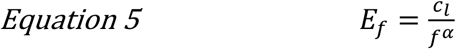

Where *c*_*l*_ is estimated contrast energy at the lowest spatial frequency, *f* is the frequency, *α* is the slope, and *E*_*f*_ is the predicted energy. *c*_*l*_ and *α* are free parameters, estimated by ordinary least squares. The function was fit to the raw data (8 spatial frequencies × 26,107 images) rather than, for example, the mean contrast of each image concept at each spatial frequency. We also fit a GLMM grouping the raw data by concept. This model leverages the organisation of images into concepts to improve the estimate of the overall model fit. The model assumes that the relationship between contrast energy and spatial frequency is distributed normally across concepts, making possible a description of the data at the individual concept level. Without a specific research question, however, this model would require extensive description of parameter estimates that is beyond the scope of this report. Importantly, the GLMM produced a group-level fit that was highly similar to the simpler linear model (1.3 for linear model versus 1.39 for multi-level model), and so we focus interpretation on only the more common, simpler “1/f” model. The 1/f model provides a similar group-level description as the GLMM but cannot be interrogated to the same extent. For example, future investigations could examine whether the random effects produced by the GLMM are sufficient to predict image concept based on spatial frequency contrast distributions alone, and so this model is included in the shared analysis code.

We fit a custom function to model contrast as a function of orientation. This model is based on a function that has been used previously to model anisotropies in perception and memory (Taylor & Bays, 2018; Wei & Stocker, 2015):

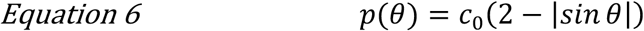

Here, *θ* is orientation in radians, *c*_0_ is a normalising constant, and *p*(*θ*) is the probability of *θ*. To capture the expected tendency that horizontal contrast is greater than vertical contrast (Hansen & Essock, 2004), we modified the function so that the distribution of energy is itself modified by a sinusoidal function centred on horizontal energy:

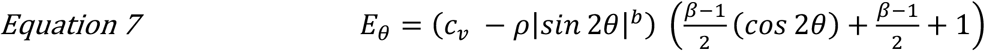

Where *c*_*v*_ is the estimated contrast for vertical orientations (i.e. when θ = ±*π*), *ρ* sets the magnitude of the difference from peak to trough, *b* sets the rate of drop-off of energy from cardinal to oblique orientations, *β* is horizontal energy as a proportion of vertical energy, and *E*_*θ*_ is the estimated energy. The parameters *c*_*v*_, *b, β*, and *ρ* are free parameters. As with the model estimating contrast energy from spatial frequency, the function was fit to the raw data (16 orientations × 26,107 images). We did not attempt this model within a GLMM framework.

### Predicting image concepts from luminance and contrast

We classified images into the concept labels supplied in the THINGS database using a simple custom linear classification analysis of the joint distributions of luminance and contrast. The general design of this analysis was motivated by Grootswagers et al (2021) who used EEG to decode which of two image concepts had been displayed to observers. In brief, we sampled every possible pair of concepts, and, for each image in each concept, we classified to which of the two concepts the image belongs using only luminance and RMS contrast. Each image, therefore, was classified according to a comparison of its own concept against every other concept one pair at a time. For example, given that there are 14 images in the first concept, “aardvark”, and 1,853 other concepts with which aardvark images were compared, there were 14 × 1853 classifications for images in the first concept alone. In total, there were 26,107 (images) × 1,853 (combinations), which is over 48 million classifications. For each pair of concepts, we found the mean proportion of correct classifications.

We first computed the mean luminance and contrast for each concept. Mean luminance is:

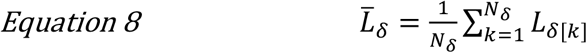

Here, *L*_*δ*[*k*]_ is the luminance of the k-th image in concept δ, *N*_δ_ is the number of images in that concept, and 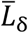 is the resulting mean luminance for that concept. Mean contrast, 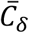, is computed similarly:

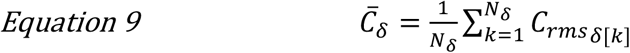

The luminance and contrast standard deviations were computed as follows:

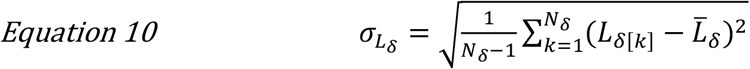

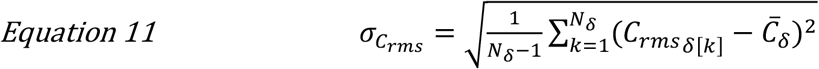

Having computed the means and standard deviations for each concept, we found the Euclidean distance of each image from every concept:

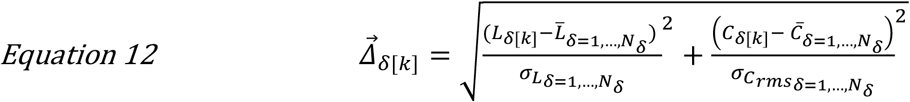

Where 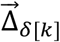 is a vector of the distances of image k from every concept, including its own. The subscript *δ*[*k*] refers to a given image (*k*) from a given concept (*δ*). Note that the distances are scaled by the standard deviation of luminance (*σ*_*L*_) and contrast 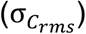 of each concept. While these distances can be used to classify images into their concepts, the results would be biased: each image’s luminance and contrast data contribute to the means of its own concept calculation, and so classification accuracy would overestimate how well *new* images could be classified. To remove this circularity, we used a leave-one-out cross validation approach in which each to-be-classified image did not contribute to the calculation of its own concept means. This analysis is equivalent to asking how accurately we could classify a held-out or unseen image into its correct concept, based only on its luminance and contrast and the luminance and contrast of the other images within each THINGS concept.

For each concept, we re-computed n-new concept means and standard deviations that excluded each image within that concept. For example, there were 14 unique images in the first concept, and so we computed 14 sets of means and standard deviations, where each set represents the means and standard deviations after leaving out a different image. Rather than compute these statistics from scratch using Equation 8 - Equation 11, it was pragmatically more efficient to compute the leave-one-out means as follows:

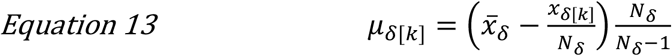

Where μ_δ[*k*]_ is the mean (either luminance or contrast) of concept *δ* after holding out image *k* from that concept. 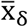 is the mean luminance or contrast of the concept with all images, while x_δ[*k*]_ is the luminance or contrast, respectively, of the held-out image. The leave-one-out standard deviations were computed as follows:

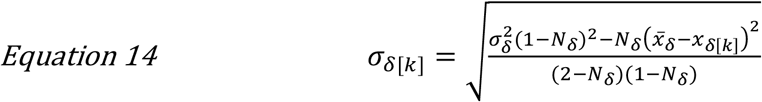

Where *σ*_*δ*[*k*]_ is the standard deviation (of either the luminance or contrast) of concept *δ* after holding out image *k* from that concept. *σ*_*δ*_ is the standard deviation of the luminance or contrast for the concept with all images. Finally, we computed the distance of an image from its own concept mean after having left out that image as per Pythagorean theorem shown in Equation 12. There were then 26,107 new distances of each image from its own concept luminance-contrast centroid. These distances were substituted into 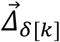, the vector calculated in Equation 12, to replace the biased estimates with left-out estimates.

The critical aspects of the equations above are that, 1) for each image, we computed the distance between each image and every concept and 2) the distance between an image and its own concept was unbiased by the image. For every possible pair of concepts, we then classified a given image from the pair as belonging to the concept that was closest to the image in luminance and contrast. If the closest concept was the image’s concept, then we scored the classification as accurate (i.e. 1); otherwise we scored the classification as incorrect (i.e. 0). Finally, we found the mean proportion correct for every pairwise concept comparison, yielding a 1,854×1,854 matrix of mean proportion correct classifications, where each matrix cell represents mean accuracy for classifying images from two competing concepts.

This simple classification model labels images into their concepts with the accuracy expected from a template-matching ideal observer who has access to only luminance and contrast data. Accuracy could be improved by using more complex models, such as a multivariate logistic regression model or a support vector machine, but these are unnecessary for our purposes of illustrating the decodability of image concepts based solely on luminance and contrast of individual images. If our simple classification model can correctly classify images into their concepts, more sophisticated models would perform better, but the interpretation would be the same: low-level images factors confound high-level category labels.

## Results

We quantified the luminance and luminance contrast in each of the 26,107 THINGS images using common metrics. All analyses were performed on grey-scaled versions of the images, but we describe how luminance modifications can be made to coloured images at the end of this section.

### Contrast energy within separable spatial frequency and orientation bands

Figure 2 shows the distributions of contrast energy as a function of spatial frequency (Figure 2A) and orientation (Figure 2B). Both sets of results are highly typical of natural images: energy scales inversely with spatial frequency (on log-log axes) and there is an over representation of cardinal orientations relative to oblique orientations. The slope of the decline of energy with increasing spatial frequency was 1.3, which is in close agreement with measurements of other natural images (Bex & Makous, 2002; Torralba & Oliva, 2003). The slope calculated by the GLMM was 1.39, with a standard deviation of 0.1 across concepts. The relatively non-overlapping distributions in Figure 2A show that the relationship between contrast energy and spatial frequency holds for all images in the THINGS database. The relatively broad overlapping distributions in Figure 2B, however, show that there is greater variation in the distribution of contrast energy over orientations across THINGS images. Regardless, we found that horizontal contrast was 1.1 times greater than vertical contrast on average (parameter *β* in Equation 7).

**Figure 2.**
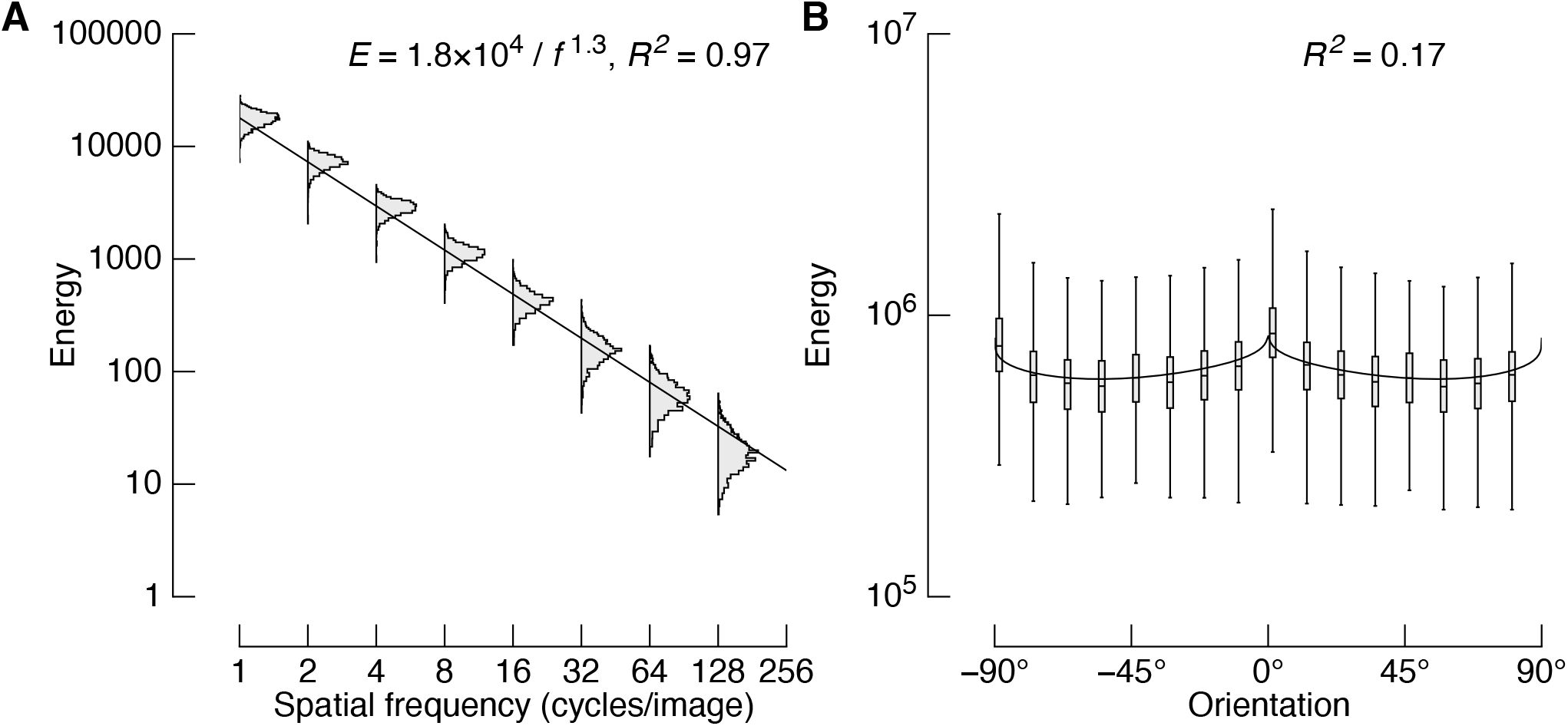
The distribution of contrast energy in separable spatial frequency and orientation bands. A) Contrast energy is inversely related to spatial frequency, following the typical 1/f^α^ relationship. Distributions are histograms of energy for all 26,107 images at each spatial frequency. The solid line is the inset function. B) Contrast energy is greater for cardinal orientations than oblique orientations. The lower and upper limits of boxes show the first and third quartiles, respectively, box centres show the medians, and the whiskers show the range. The solid line is a model that captures the cardinal/oblique biases, as well as differences in horizontal versus vertical contrast, as described in the Methods.

### Relationship between luminance and RMS contrast

Figure 3A plots the luminance and RMS contrast of every image in the THINGS database. The inset marginal distributions show that both measures are approximately normally distributed. Luminance tends to be lower than mid-grey, and RMS contrast tends to be distributed around .5. The line through the data is the best fitting quadratic model fit in a GLMM framework (Equation 4). The same data are plotted in Figure 3B, with three randomly drawn concepts highlighted in different colours. Note that the means of the concepts are separable in both luminance and contrast, and there is partial overlap of the distribution of images from different concepts. These data suggest that these simple image statistics may be sufficient to classify images in their concepts with above-chance accuracy.

**Figure 3.**
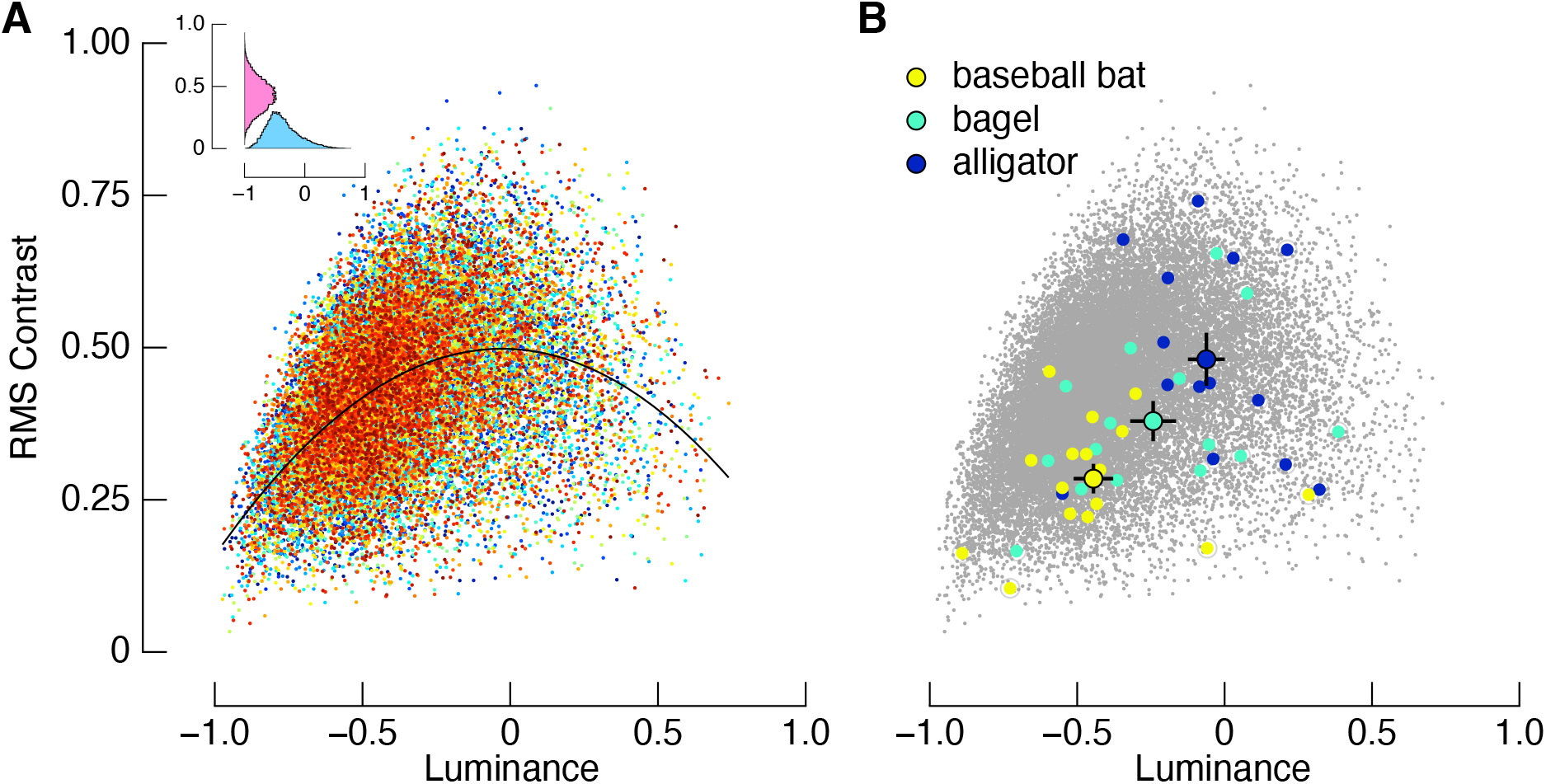
Relative luminance and RMS contrast of 26,107 images in the THINGS database. A) The relationship between luminance and RMS contrast is parabolic, such that there is a positive correlation for negative luminance images, and a negative correlation with positive luminance images. Each datum has been colour coded according to image concept but note that there are more image concepts than colour categories. The inset at top left shows the marginal histograms. Relative to mid-grey, images have a lower luminance on average (blue distribution), and are approximately normally distributed around 0.5 RMS contrast (pink distribution). B) Measures for three randomly selected image concepts, and their averages. Average measurements are shown in black outlines, with error bars showing one standard error. See Methods for details on scaling and calculations.

### Concept classification based on luminance and contrast

We next tested how accurately each image could be classified into its concept using luminance and contrast alone. This analysis is motivated by the fact that some variables of theoretical interest will be correlated with changes in luminance and contrast, confounding some experimental designs that leverage the THINGS images as stimuli. Such dependent variables include changes in BOLD or the amplitude of an event-related potential in neuroimaging experiments. We discuss these issues more in the Discussion.

Classification accuracy for all 1854^2^ pairwise comparisons are shown in Figure ***4***4A. We consider classification accuracy as being analogous to representational dissimilarity measures (Grootswagers et al., 2021): the greater the decoding accuracy of a pair of concepts, the more dissimilar they are in luminance-contrast space (e.g. Figure 3B). 63.4% of all images were correctly classified, compared with chance classification of 50%. The absolute accuracy of decoding is less important than the fact that accuracy is well above chance. This result shows that low-level image factors are a source of decodable information. Critically, these low-level factors are confounded with the high-level concept labels of the THINGS images.

**Figure 4.**
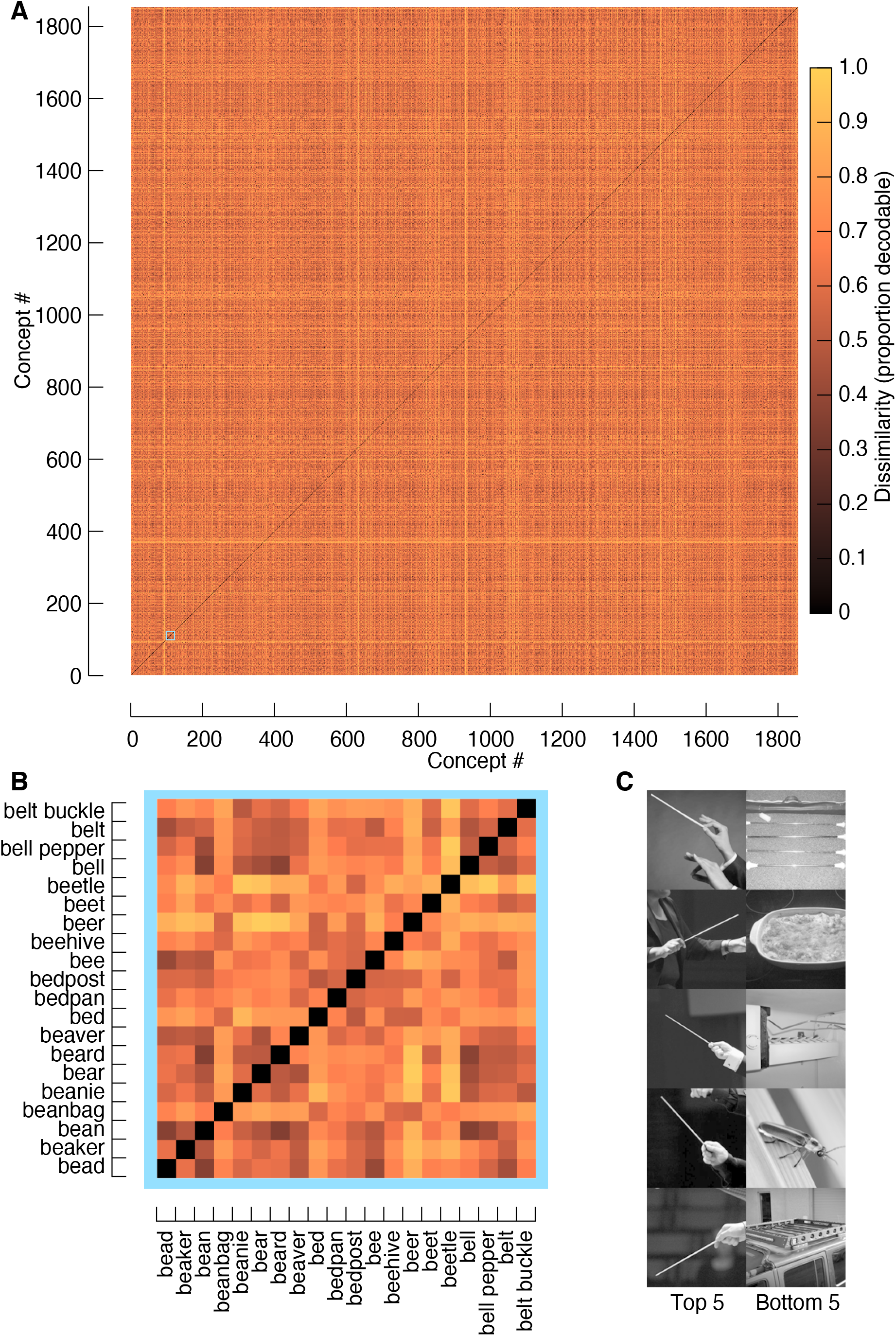
THINGS images can be classified based on their mean luminance and RMS contrast alone. A) Proportion of correct concept discriminations for all possible pairwise comparisons of concepts. Within each image concept, there were at least 12 image exemplars that were classified as belonging to one of the 1,854 concepts. An image was classified by minimising its luminance and RMS difference from each concept average. Chance classification is 0.5. B) A detailed section of (A), highlighting the classification accuracy for a subset of 20×20 concept pairs. C) Five images that were most and least accurately identified.

Figure ***4***4B shows a zoomed in section of the dissimilarity matrix with concept labels. As one example of differences in classification accuracy of images, when paired with beetle images, beanie images can be classified more accurately than belt buckle images. The five most and least accurately classified images are shown in Figure 3C. While the top five most classifiable images were from the same concept (“baton”), the bottom five classifiable images come from different concepts. These example images provide an intuition behind the (un)successful classification of images. The top five images clearly differ from the bottom five in both their mean luminance and pixel variability (i.e. RMS contrast). Image concepts that cluster near the centroid of the distribution in Figure 3A will have more similar image metrics than image concepts near the edges of the distribution, and therefore will be less discriminable based on these metrics alone.

The above-chance level of classification for paired concepts, described above, motivated us to test how well we could classify each image into its concept when compared with every other concept within the database. This analysis was the same as used for pairwise classifications, but we classified a given image according to the smallest distance between the image’s statistics and the nearest of all concept means. In this case, chance accuracy is 1/1854, or 0.05%. Using luminance and contrast alone, mean classification accuracy was 0.3% (79 images out of 26,107 were correctly classified). Although only a small minority of images were correctly classified, this level of accuracy is significantly greater than chance (binomial test: Successes = 79, N = 26107, P = 1/1854, p < 10^−10^). We repeated the classification analysis, now including the spatial frequency and orientation contrast energy data presented in Figure 2 in addition to luminance and RMS contrast. By including these predictors, classification accuracy increased to 1.5% (400 images out of 26,107 were correctly classified; p < 10^−10^).

## Discussion

In the present report we summarise the luminance and luminance contrast properties of images in the THINGS database. There were three primary results. First, the distributions of contrast energy over spatial frequency and orientation were similar to the statistics reported for other natural images (Figure 2). Second, there was a nonlinear but predictable relationship between mean luminance and RMS contrast across images (Figure 3). Third, we showed that there are systematic differences in luminance and contrast between image concepts that are sufficient to classify individual images into their concepts with above-chance accuracy (Figure ***4***4). In the following, we discuss the implications of these findings in more detail, and provide an example of how to normalise luminance and contrast of coloured images to remove these factors as potential experimental confounds. We also discuss the importance of natural image databases, like THINGS, for advancing cognitive neuroscience theory.

### Contrast energy

The contrast energy of THINGS images was well described as a linear transform of spatial frequency on log-log axes, with a slope of 1.3 (Figure 2A). There was relatively little variation across images, with the simple 1/f model accounting for 97% of variance across all 26,107 images. Contrast energy as a function of orientation, however, was relatively more heterogeneously distributed across images (Figure 2B). Nonetheless, our model of oriented contrast accounts for 17% of the variance in the THINGS images. There are many plausible reasons for the greater consistency of energy as a function of spatial frequency relative to orientation. As one example, Torralba and Oliva (2003) showed that such statistics depend on whether image content is natural or not, with relatively less cardinal-oblique contrast bias for natural images. A strength of the THINGS database is its diversity of image content. Furthermore, by design, the images in the THINGS database are digital photos with objects in various compositions and configurations. For example, some photos tightly frame a specific object, and others are framed with the camera lens rotated by some amount. These variations will have little impact on the 1/f spectrum of the images, which simply quantifies the scale invariance of the contrast distributions, but can greatly impact the distribution of oriented contrast (e.g. a picket fence rotated by 45° will have the same 1/f spectrum as the original, but its oriented contrast distribution will be translated). It is somewhat unsurprising, therefore, that oriented contrast is more variable across images. Perhaps more important to the intended use of THINGS images, however, is that there is no reason to expect that these distributions of contrast energy deviate meaningfully from those encountered in natural settings.

### Luminance and RMS contrast

Mean luminance and RMS contrast were correlated according to a quadratic function: for images with negative mean luminance, there is a positive relationship between luminance and RMS contrast; for images with positive mean luminance, there is a negative relationship between luminance and RMS contrast. The likely explanation for this nonlinear relationship is somewhat trivial: when the mean luminance of an image approaches extreme values, the distributions of pixels become skewed, resulting in lower standard deviations. Frazor and Geisler (2006) used calibrated photos to estimate the luminance of images as they would have appeared in the world (as opposed to their luminance on a computer monitor), and found no systematic relationship between luminance and RMS contrast.

A clear potential benefit of the THINGS database is to facilitate understanding of how high-level conceptual information influences perceptual, cognitive, and neural representations of visual input (Grootswagers et al., 2021; Harrison, 2019; Hebart et al., 2019, 2020; Neri, 2011; Rideaux et al., 2021). However, the variation in luminance and contrast in the THINGS images may be an undesirable property for studies in which these factors directly influence measures of interest. Consider a hypothetical experiment that uses pupil diameter as a dependent variable. A researcher may ask whether pupil diameter predicts the frequency with which THINGS concepts appear in a given text corpus. Pupil diameter will be responsive to the luminance and contrast of each image, and we have now shown that image concepts can be predicted from these same factors alone. Therefore, the hypothetical experiment predicting concept frequency from pupil diameter would be confounded by low-level image factors unrelated to concepts per se. This basic confound would also need to be ruled out of neuroimaging experiments in which event related potentials or changes in BOLD signal depend on luminance and contrast.

Whereas many recent models of image classification are deep neural networks (e.g. Hebart et al., 2019), we showed that a highly simple model is sufficient to achieve well above chance classification accuracy. Deep neural nets provide a means by which one can find low-dimensional structure in natural images that is predictive of image content. By contrast, our model uses only luminance and RMS contrast information to classify images into their concepts. We classified an image by finding the concept whose mean luminance and RMS contrast were closest to the image (see Figure 3B and Figure ***4***4 and Methods). We achieved 63.4% accuracy when only two concepts were used as per Grootswagers et al (2021). High-level image structure, therefore, is not necessary for above-chance classification of high-level information. The ability to classify images using luminance and contrast alone demonstrates the potential impact these quantities could have on physiological responses to, and the neural correlates of, the THINGS images.

### Normalising luminance and RMS contrast across images

There are simple ways to guard against differences in luminance and RMS contrast confounding experiments that leverage THINGS images as stimuli. In Figure 5 we show an example in which we normalise the luminance and RMS contrast of the full colour images. The basic process involves converting the RGB images into a format in which luminance is dissociated from colour. We use YUV, in which the matrix Y contains the image’s luminance (Figure 5A). Normalisation involves simply subtracting the mean luminance and dividing the result by the standard deviation. The resulting luminance plane can be multiplied by some constant that represents the desired RMS contrast (0.23 in this case). By recombining the luminance and colour planes, the image can then be converted back into RGB space. After this normalisation process, images are matched in their luminance and RMS contrast. This process effectively reduces the distribution of luminance and RMS contrast across all images in Figure 3 to a single point. Note that, although the images may tend to appear qualitatively faded, the objects (and concepts) remain clearly visible in the normalised images (Figure 5B). Applying the same decoder as described in the Methods and Results to these normalised images reduces decoding accuracy to 49.9%. The simple normalisation method presented here, therefore, removes luminance and contrast as decodable sources of information.

**Figure 5.**
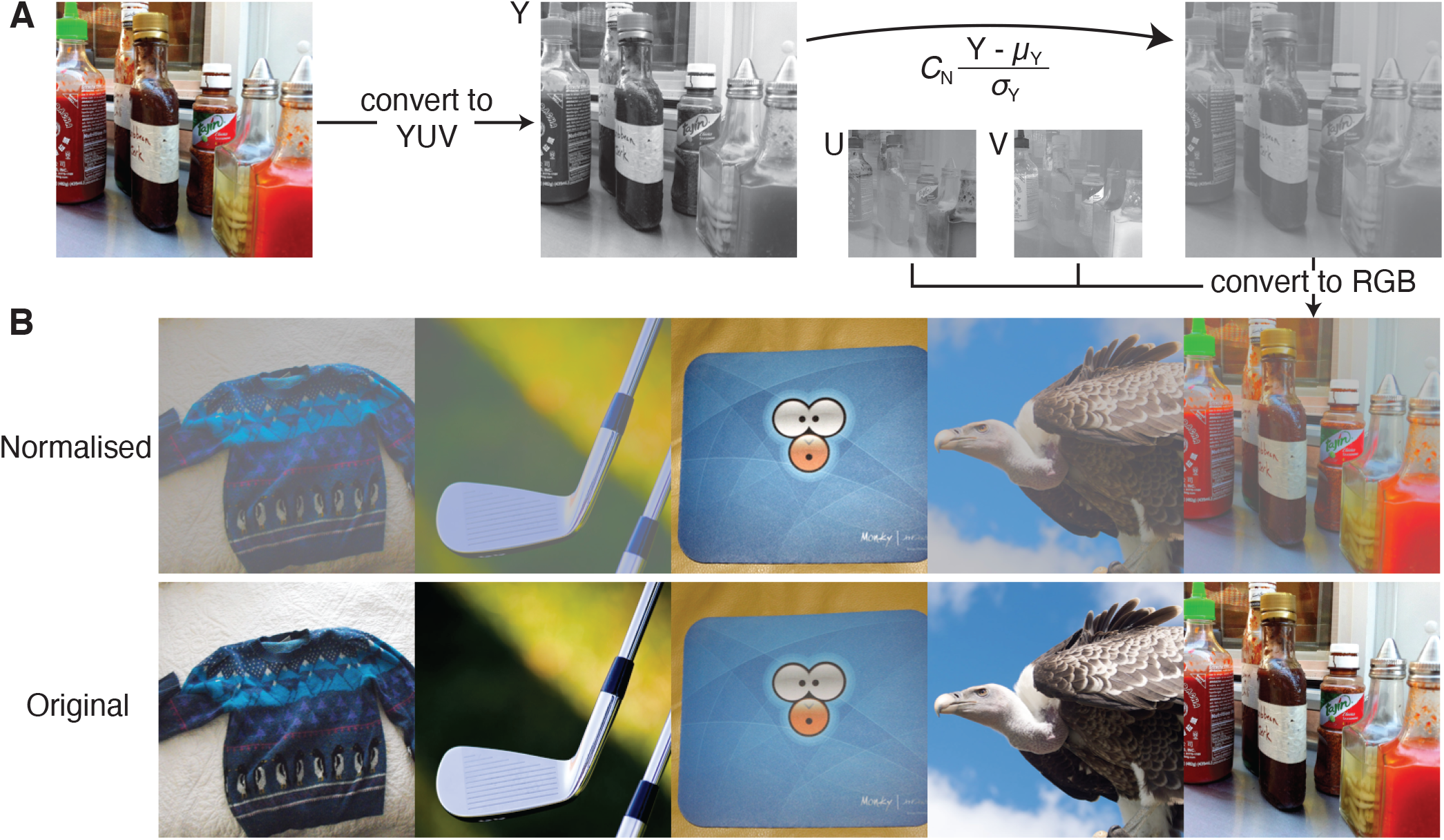
Example procedure to normalise luminance and contrast. A) We convert an RGB image into YUV, where Y is the luminance of the image. Y is then normalised by removing its mean, μ_Y_, and dividing by its standard deviation, σ_Y_. RMS contrast can be set by a multiplicative constant, C_N_. By combining this new luminance plane with the original UV images, the image can be converted back to an RGB image, now with a mean luminance of zero and RMS contrast of C_N_. B) Examples of normalised images (top) compared with their originals (bottom). All images in the top row have the same mean luminance (0) and RMS contrast (0.23).

### Other stimulus considerations

When we included in our decoding analysis the contrast energy within spatial frequency and orientation bands in addition to RMS contrast and mean luminance, decoding accuracy improved five-fold (i.e. from 0.3% to 1.5% accuracy at classifying each image into one of 1854 concepts). This improvement comes from increasing the dimensionality of the summary data; the decoder had access to more information about each image. It may therefore be pertinent for researchers to also control for these factors. Indeed, Willenbockel et al (2010) reviewed several neuroimaging studies in which such basic low-level image factors may have influenced results. They developed and made available the SHINE toolbox to control for a broader range of image factors than those reported here. We refer readers to that useful resource for both a more detailed review of potential confounds in previous neuroimaging studies as well as their software. We note, however, that when using natural image stimuli, image factors *must* differ between images. It is up to the researcher to determine which image statistics they are interested in, and which ought to be matched across images. As shown in Figure 5B, images that have had their image statistics manipulated remain easily recognisable; human image recognition is invariant over a large range of mean luminances and contrasts. We suggest that researchers interested in relatively high-level image features adopt a conservative approach to these issues and assume that, if low-level image factors *could* contaminate their models, then they probably do. This holds for behavioural, neural and machine-learning models.

Various image metrics will be affected by decisions such as the display size of images and how an image is converted to greyscale from colour space (and back again). We converted to greyscale because we were interested only in luminance and luminance contrast. We resized images because it was more convenient to calculate image metrics from a constant-sized input. When increasing or decreasing the size of an image, one has to make a choice about how to add or remove pixel information, respectively. In our tests, using an interpolation method to either increase or reduce image size holds the mean (luminance) constant while reducing the variability (RMS contrast). Without interpolation, the mean is more variable when resizing the image, while the variability remains relatively constant. Perhaps most critical for the application of this knowledge, however, is that any calculations or controls that are performed on stimuli must be performed on the image as it will be (or was) presented to the observer.

There are, of course, good alternatives to adjusting the low-level features of stimuli to mitigate confounds in experiments. Depending on the research question and experimental design, simply inverting the image may be sufficient to control low-level factors while disrupting higher-level and semantic processing (e.g. Neri, 2014; Yin, 1969). In other cases, the same stimuli may be repeated in all conditions, while the experimenter manipulates only the instructions given to participants across conditions (e.g. Harrison et al., 2019; Harrison & Rideaux, 2019).

### The THINGS database is a valuable resource

We do not believe that any of the results presented here should discourage researchers from using THINGS images as experimental stimuli. Instead, we encourage the use of the THINGS database. The use of natural images as experimental stimuli will help to bridge the gap between the minimal stimuli typically used in studies of cognitive neuroscience and the real world. We hope our data will facilitate a better understanding of basic statistical properties that may be important to future research. The close match of the image spectra to known natural image statistics demonstrates that these images do not deviate from other natural images in a meaningful way. While we found that systematic luminance and contrast differences across image concepts have the potential to confound experimental designs, these properties can be easily adjusted. Adjustments can be made a priori by, for example, normalisation, or post-hoc, by using our shared data to partial out variance attributable to these low-level statistics independently of other experimental manipulations. One improvement for future iterations of either the THINGS database or other similar resources would be to include any available metadata regarding the calibration, brand and model of digital camera from which an image was obtained in order to correctly linearise images prior to analysis and use in experiments.

### The importance of natural images in cognitive neuroscience

Much of our knowledge of visual perception and cognition comes from experiments with relatively abstract stimuli. The functions of neurons in primary visual cortex are some of the best understood in cognitive neuroscience. Much of this understanding has been derived from oriented gratings of various spatial frequency, contrast, and position in the visual field (for a review, see Carandini et al., 2005). There is evidence that such knowledge, derived from experiments with minimal stimuli, is sufficient to predict neural encoding of complex natural images (Cadena et al., 2019; Kay et al., 2008; Nishimoto et al., 2011; Sebastian et al., 2017; Yoshida & Ohki, 2020). In many other areas, however, there seems to be little emphasis on generalising theory beyond the abstract stimuli arbitrarily chosen by the experimenter. This seems particularly true of the cognitive neuroscience of visual attention and working memory – we refer readers to a review chapter by Brady and colleagues (2019) who discuss this issue at length. Whether a theory is useful for more than predicting how volunteers respond to spots of light on a computer monitor requires bringing experiments out of the lab and into the real world. Such a leap will likely come at the cost of experimental control. However, large stimulus datasets like the THINGS images provide a steppingstone between standard experimental stimuli and the complexity of the real world. THINGS images are highly diverse in their content and capture much of the diversity of natural scenes, but, as shown here, can nonetheless be described and controlled in terms of their low-level properties.

Are digital photos sufficient to understand perceptual and cognitive processes when viewing “true” (non-digital) natural images? Anecdotally, a common criticism levelled at investigations that treat digital photos as representative of natural images is that the images are photographed for a specific purpose by a specific photographer, and therefore may not truly represent the visual images we encounter in nature (see Hart et al., 2009 for one example in which lab-based experiments are directly compared with the real world). Although there may remain a distinction between processes involved in viewing digital photos versus non-digital images, we do not think the importance of digital photos in psychology experiments is undermined for at least three reasons. First, digital photos provide a justifiable balance between uncontrolled complexity and experimental control, and this balance requires more powerful and generalisable theory (Carandini et al., 2005; Muthukrishna & Henrich, 2019). Second, the human visual system is putatively shaped by the *statistics* of natural environments over evolutionary and developmental timescales, and, in general, there is no reason to expect that the statistics of digital photos differ meaningfully from the statistics of natural environments (Geisler, 2008; Olshausen & Field, 1996). Third, Hebart et al (2020) have shown how digital photos and perceptual decisions can be combined to create data-driven models that generate meaningful psychological constructs that generalise across images and contexts. Any limitations of digital photo stimuli also do not justify the sole use of arbitrary abstract stimuli in cognitive neuroscience.

## Summary and conclusions

Our analyses show that the THINGS images have typical distributions of luminance and contrast, and that these low-level image factors can be used to classify individual images into their concept. The separability of luminance and contrast may confound certain experimental designs, but can be controlled for easily. We see great potential in the THINGS database as a large diverse stimulus set that can be used to address a great variety of research questions in cognitive neuroscience. We are already exploiting this resource to understand perceptual processes in natural images (e.g. Rideaux et al., 2021). Wide adoption of the database will likely result in yet even larger open datasets when researchers make available their results. We anticipate that the cumulative massive datasets will create new and unique opportunities to link previously disconnected cognitive neuroscience theory via a common stimulus set. This trans-disciplinary information is already being curated: https://things-initiative.org/

## Acknowledgements

I am grateful to Tijl Grootswagers who provided feedback on an earlier draft of this manuscript and explained to me the importance of using a cross-validation analysis for image classification. I would not have done this step of the analysis without his feedback and guidance. I also thank Cameron Turner who worked out how to calculate the leave-one-out standard deviations shown in Equation 14. This research was supported by an Australian Research Council Discovery Early Career Research Award (DE190100136).

